# Nuclear blebs are rupture-prone restricted compartments of the nucleus

**DOI:** 10.64898/2026.11.15.766853

**Authors:** Jimena Collado, Mai Pho, Marilena L. Currey, Katherine Chiu, Cameron Herman, Yasmin Berrada, Vadim Backman, Luay M. Almassalha, Andrew D. Stephens

## Abstract

Nuclear blebs are abnormal herniations of the cell nucleus that cause nuclear rupture and dysfunction. The biophysical principles underlying nuclear blebs and ruptures remain to be fully elucidated. Using nuclear localization signal green fluorescent protein (NLS-GFP) we tracked nuclear bleb rupture dynamics. Upon nuclear rupture, the nuclear bleb’s NLS-GFP loss rate is significantly faster than the nuclear body, whereas non-blebbed nuclei ruptures show no difference in NLS-GFP loss rates between the site of nuclear rupture and body.

Micromanipulation compression induced nuclear bleb rupture recapitulates 5-fold nuclear bleb loss rate. Furthermore, fluorescence recovery after photobleaching (FRAP) in the intact nuclear bleb is significantly slower than recovery in the nuclear body. Confocal imaging reveals that the transition from the nuclear bleb to body is significantly constrained. Thus, we reveal the nuclear bleb as a restricted compartment and nuclear rupture site, both of which greatly affect nuclear functions.

## Introduction

The nucleus is the organelle that holds and compartmentalizes the genome and the vital biochemical processes that control reading of the genome which ultimately determines the bulk of cellular behavior. The nucleus and its main mechanical components chromatin and lamins must properly resist cytoskeletal and/or external forces to maintain the integrity of the nucleus as a compartment and its shape (Stephens *et al*., 2019a; Kalukula *et al*., 2022). Abnormal nuclear morphology is a major hallmark of human disease spanning aging to heart disease to cancer and many others. A specific type of abnormal nuclear morphology is called nuclear blebbing, in which the interphase nucleus forms a herniation that is greater than 1 µm and has decreased DNA density (Bunner *et al*., 2024; Chu *et al*., 2025; Pujadas Liwag *et al*., 2025; Clark *et al*., 2026). Nuclear blebbing causes nuclear rupture, the loss of nucleus compartmentalization resulting in spilling of nuclear contents into the cytoplasm and vice versa. Nuclear blebbing and rupture is well documented to lead to cellular dysfunction by causing disrupted DNA damage and repair (Chen *et al*., 2018; Xia *et al*., 2018; Shah *et al*., 2021; Pho *et al*., 2023), transcription (De Vos *et al*., 2011; Helfand *et al*., 2012; Berg *et al*., 2023), and cell cycle control (Pfeifer *et al*., 2018). Alterations in these core cellular functions of DNA damage and repair, transcription, and cell cycle can themselves increase nuclear blebbing and rupture (Eskndir *et al*., 2025; Bunner *et al*., 2026; Prince *et al*., 2026). Nuclear blebs are plastic deformations that remain for long periods of time and require cell division to reset nuclear shape (Zheng *et al*., 2026). This nuclear rupture and dysfunction feedback could cause an ever-worsening diseases state. However, physical underpinnings of nuclear blebs and ruptures remain unclear.

Recent work has provided key metrics to understand the physical basis for how blebs and ruptures drive cellular dysfunction. One study measured the micropipette aspiration suction pressure required to induce different levels of nuclear rupture, measured to percentage decrease of NLS-GFP loss from the nucleus (Zhang *et al*., 2019). Many other studies have outlined how induce nuclear blebbing and rupture through perturbations (Denais *et al*., 2016; Raab *et al*., 2016; Stephens *et al*., 2018, 2019b) or through new tools such as micromanipulation compression (Currey *et al*., 2022). Other studies have outlined changes in nuclear height via confocal imaging providing key measurements in 3D to provide understanding how the nuclear body and bleb sizes differ (Hatch and Hetzer, 2016; Pho *et al*., 2023).

More recently studies have reported that nuclear blebbing, independent of nuclear rupture, is associated with nuclear dysfunction (Helfand *et al*., 2012; Shah *et al*., 2021; Chu *et al*., 2025). Nuclei presenting a nuclear bleb show increased DNA damage. This is most striking in prostate cancer cell line LNCaP, which shows infrequent nuclear rupture, but still exhibits higher levels of DNA damage in blebbed nuclei (Chu *et al*., 2025) as well as transcriptional repression (Helfand *et al*., 2012). This suggests that the physical basis of the nuclear bleb may disrupt normal processes. Thus, we hypothesize that the nuclear bleb is a restricted compartment of the nucleus.

Past studies have focused on the repair of nuclear ruptures and ruptures in confined migration. However, the dynamics of nuclear ruptures in non-migrating cells remain relatively unknown. Here we image mouse embryonic fibroblasts with NLS-GFP to track nuclear shape and nuclear rupture across wild type and in three different perturbations of chromatin decompaction via histone deacetylase inhibitor (VPA), lamin B1 null cells *(Lmnb1-/-*), and lamin A/C null cells (*Lmna-/-*). Tracking and analysis of NLS-GFP loss-spill dynamics during nuclear ruptures reveals that nuclear blebs loose nuclear content faster than the nuclear body. Furthermore, fluorescence recovery after photobleaching of the bleb and body provides differential dynamics. Finally, measurements of nuclear height reveal a drastic constriction in the connection between the nuclear bleb and the body. Thus, data reveal that the nuclear bleb is a restricted compartment from the nuclear body.

## Results

### Nuclear rupture loss dynamics are 10-fold faster than nuclear envelope breakdown

To determine the dynamics of nuclear rupture, we measured the NLS-GFP loss graph of both the nuclear bleb and the nucleus during rupture. To capture the dynamics of nuclear rupture we imaged NLS-GFP cells over 45 minutes at 5 second intervals for four different conditions. We imaged and analyzed bleb-based nuclear ruptures in wild type, chromatin decompaction perturbation (VPA), and loss of lamin B1(*Lmnb1-/-*), in which bleb-based ruptures are the majority form of nuclear rupture (Pho *et al*., 2023). We also imaged and analyzed non-bleb-based nuclear ruptures which occur as the major form of nuclear rupture in lamin A/C null cells (*Lmna-/-*) for comparison.

To better understand the speed of loss of NLS-GFP upon nuclear ruptures we compared it to nuclear loss of compartmentalization in mitosis of wild type cells (**Figure 1B**). The nuclear envelope breakdown upon entry into mitosis is a control for loss of compartmentalization (Beaudouin *et al*., 2002). Loss of NLS-GFP during nuclear envelope breakdown before mitosis shows a 10-fold slower loss rate than compared to nuclear ruptures (∼10% loss over 100 seconds for mitosis vs ∼10% loss in 5 seconds post-rupture; **Figure 1B**). We also note that nuclear ruptures plateau within 30 seconds at a drastic NLS-GPF loss of 30-50%. While we image at small time intervals, overall the nuclear body NLS-GFP loss rates appear similar to other published reports with longer time intervals (Vargas *et al*., 2012; Young *et al*., 2020; Pho *et al*., 2023). Thus, this fast, temporary, and drastic loss NLS-GFP suggests nuclear ruptures are a high- pressure short-lived event.

**Figure 1.**
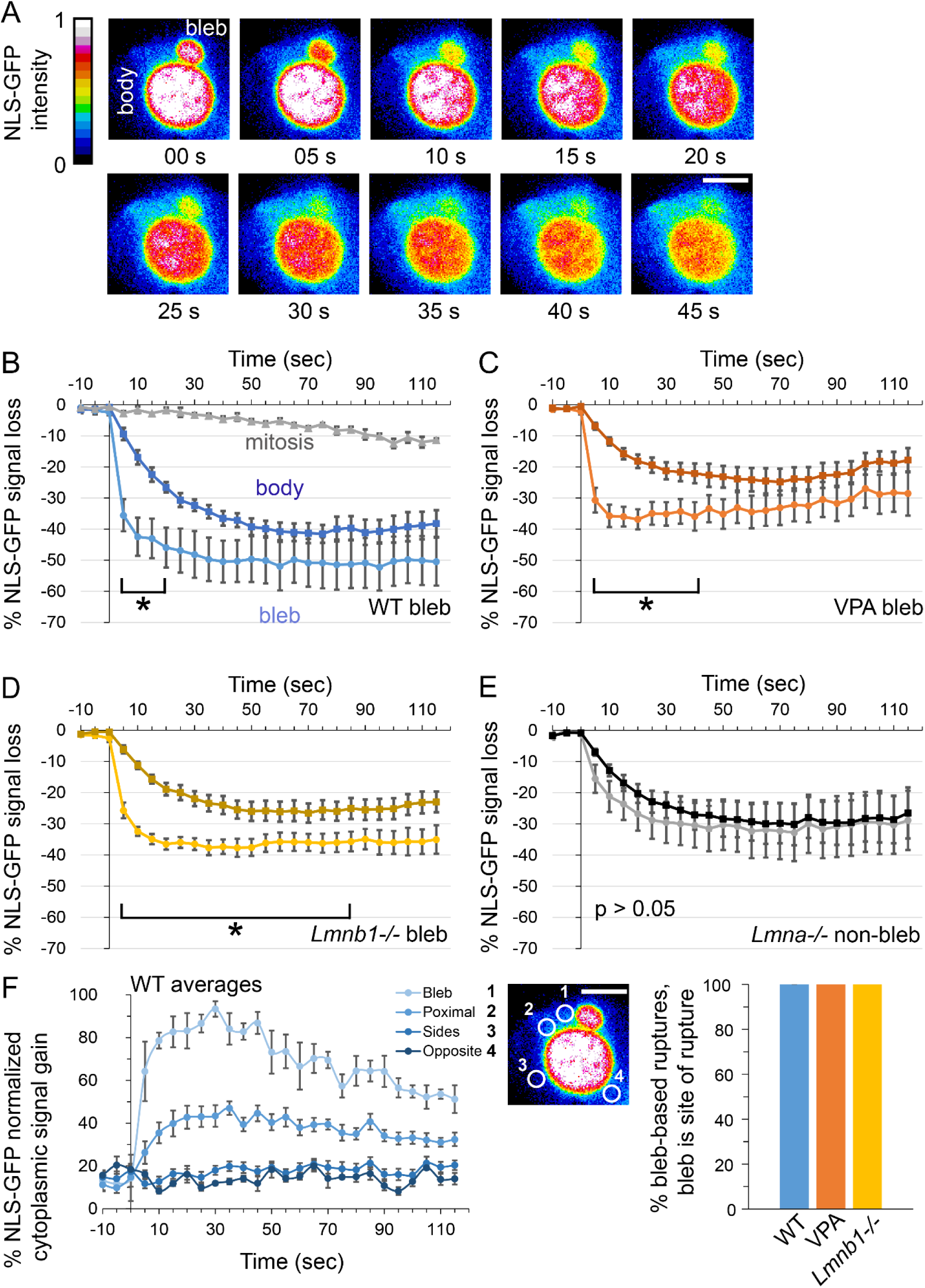
Nuclear rupture dynamics reveal that the bleb is a separate compartment and the consistent site of rupture. (A) Example timelapse of a blebbed nucleus during a rupture event displayed in color scheme representing NLS-GFP intensity. (B-D) Average intensity of the main nucleus body (dark color) vs. bleb during bleb-based nuclear rupture for WT (B, blue, n = 5), VPA (C, orange, n = 15), and *Lmnb1-/-* (D, yellow, n = 13). In B we also measured loss of NLS-GFP during nuclear envelope breakdown associated with mitosis in WT, which is displays a relatively order of magnitude slower loss rate (n = 3 cells). (F) Non-bleb-based nuclear rupture for *Lmna-/-* for rupture site (gray) vs distal body (black), n = 6. (H) Line graph of normalized to maximum NLS-GFP signal increase upon rupture at different sites, with an example image of a nucleus (same from A) showing regions of interest correspond to the bleb. Bar graph shows % of measured ruptures that occurred at the bleb (WT n = 5; VPA n = 15; *Lmnb1-/-* n =14). An asterisk denotes significantly different, p < 0.05 determined by paired Students’ t-test. Error bars represent standard error. Scaler bar = 10 µm.

### Nuclear blebs are the site of rupture and lose diffusible content faster than the nuclear body

We hypothesized that the nuclear bleb could be a restricted compartment of the nucleus. To determine if the nuclear bleb is a restricted compartment, we tracked NLS-GFP loss rates of the nuclear bleb and compared them relative to the nuclear body. If the nuclear bleb is not a restricted compartment, then loss rates upon rupture should be statistically similar. Whereas, a restriction would cause NLS-GFP loss rates to be statistically different between the nuclear bleb and body.

All measured bleb-based nuclear ruptures showed a clear qualitative pattern of the nuclear bleb losing NLS-GFP signal faster than nuclear body (**Figure 1A**). We quantify and compare nuclear bleb vs. body loss averages for 5 seconds intervals for wild type, VPA, and *Lmnb1-/-* (**Figure 1 B-E**). The data clearly show that immediately post rupture NLS-GFP loss is faster in the nuclear bleb (∼30%) than in the nuclear body (∼10%) on average for all conditions. The time interval for difference in NLS-GFP loss percentage between bleb and body varies between conditions. Thus, nuclear blebs display a reproducible faster rate of NLS-GFP loss relative to the nuclear body which suggests that there is a restriction between the nuclear body and bleb that slows its NLS- GFP loss.

To determine if this differential NLS-GFP loss rate of the nuclear bleb vs. body is simply due to the bleb being proximal to the rupture site, we measured nuclear loss rates proximal and distal in non-bleb-based nuclear ruptures which are the majority of rupture events in L*mna-/-* (Pho *et al*., 2023). *Lmna-/-* displays majority non-bleb-based nuclear rupture in which the NLS-GFP loss over time was not statistically different between the site of rupture and the distal nuclear body (**Figure 1E**). Wild type non-blebbed based ruptures also showed similar NLS-GFP loss rates for site of rupture and distal location, except for 5-second time point post-rupture which was statistically significantly different (**Supplemental Figure 1A**). These findings agree with the idea that the same compartment of the nucleus should have the same loss dynamics. Furthermore, we tracked rare nuclear blebs and their rupture in *Lmna-/-*, which recapitulated the same faster NLS-GFP loss in the nuclear bleb than the body (**Supplemental Figure 1B-C**). Thus, non-bleb-based nuclear ruptures display similar NLS-GFP loss dynamics while the nuclear bleb in bleb-based nuclear ruptures displays a significantly faster loss rate.

To confirm that nuclear blebs are the site of rupture, we tracked NLS-GFP increase in the cytoplasm to determine where NLS-GFP was leaving the nucleus. We drew 1 µm sized regions of interest around the nucleus in the cytoplasm to track where and how the NLS-GFP spills from the nucleus. We were able to locate the exact location of ruptures which show a strong local increase in NLS-GFP signal at the site of rupture which falls off with distance (**Figure 1F**). From this analysis we find that the nuclear bleb was the site of rupture for all measured events. This data nicely agrees with loss of lamin B1 only at nuclear blebs upon rupture and recruitment of repair proteins to the bleb post-rupture (Chu *et al*., 2025). Overall, nuclear blebs display a faster and different NLS-GFP loss rate than the body and are the consistent site of rupture.

### Micromanipulation induced rupture recapitulates differential compartment loss dynamics

Micromanipulation of micropipettes provides the ability to physically manipulate the nucleus (Currey *et al*., 2022). We previously showed that micromanipulation via moving point compression can induce nuclear ruptures from blebbed nuclei and even separately induce nuclear blebs. To capture NLS-GFP nuclear ruptures with higher temporal sampling we used micromanipulation and live cell imaging at 15 frames per second to induce, capture, and analyze nuclear rupture dynamics with finer details (**Figure 2A**).

**Figure 2.**
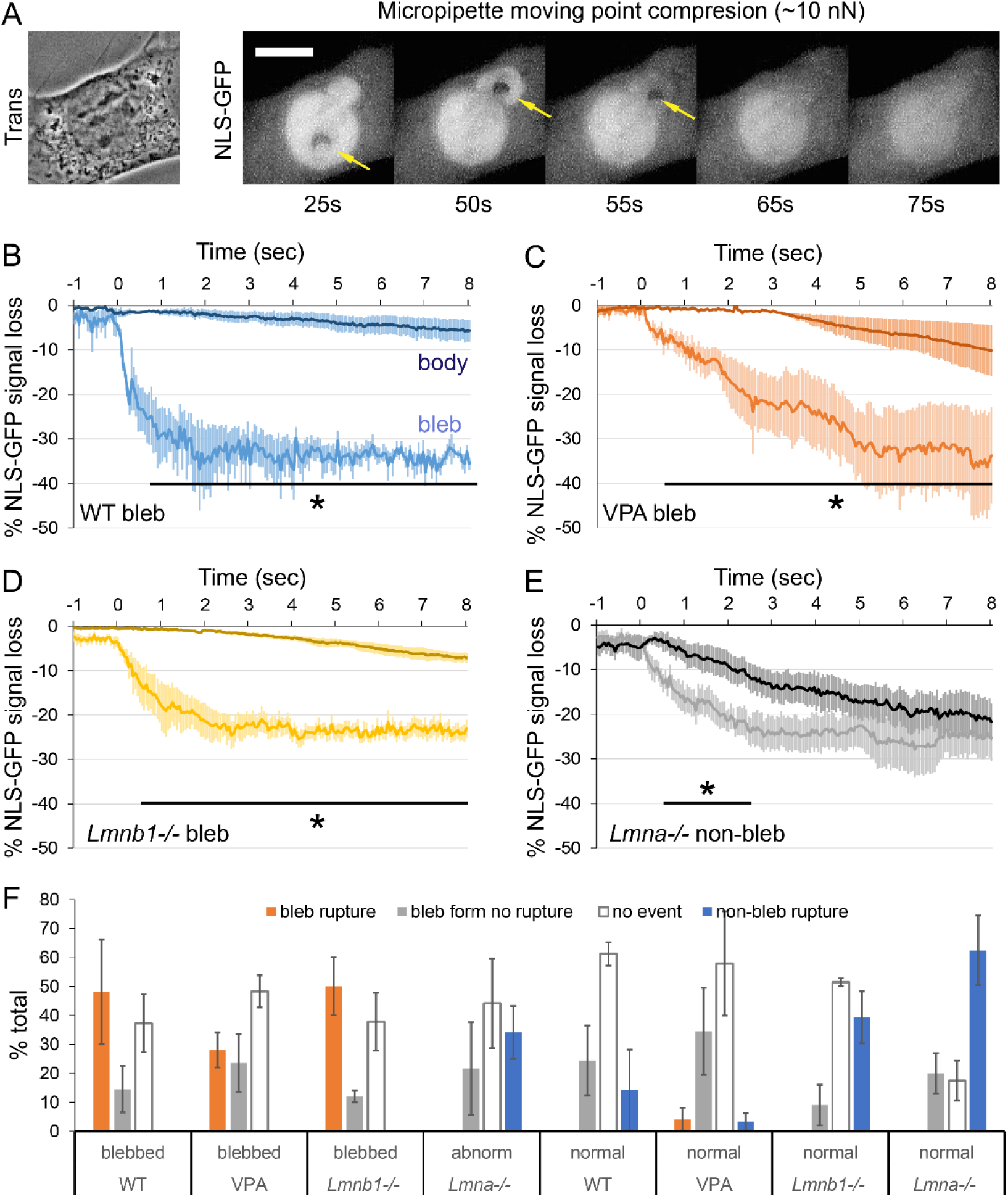
Micromanipulation local compression induces nuclear ruptures that recapitulate differential dynamics between nuclear bleb and body. (A) Images of a micropipette moving point compression experiment that induced rupture of a blebbed nucleus visualized by NLS-GFP. (B-E) Sub second (15 frames per second) tracking of NLS-GFP loss from the main nucleus body (darker color) vs. loss from the nuclear bleb (lighter color) for bleb- based nuclear ruptures of wild type (WT, B), chromatin decompaction (VPA, C), lamin B1 loss (L*mnb1-/-*, D) and non-bleb rupture in lamin A/C loss (*Lmna-/-*, E). The line represents the average loss and the emanating lighter color represents the standard error (n = 3 WT, 3 VPA, 5 LMNB1-/-, 5 LMNA-/-). An asterisk denotes significantly different, p < 0.05 determined by paired Students’ t-test. Error bars represent standard error. (F) WT, normal (7,3,5) blebbed (4,5,8,6); VPA, normal (9,10,8) blebbed (5,8,9,8); LMNB1-/-, normal (10,8,11) blebbed (9,10,11,11); LMNA-/-, normal (10,6,10,6) abnormal (10,6,6,6).

Live cell imaging of naturally occurring bleb-based nuclear ruptures displayed the majority of NLS- GFP is loss occurs in the first 5 seconds (**Figure 1**). Using micromanipulation to cause moving point compression across the nucleus to the bleb provided a manner to induce nuclear rupture and capture this initial loss rate with high temporal resolution. Similar to previous measurements, NLS-GFP loss rate was visibly faster in the nuclear bleb relative to the nuclear body (**Figure 2A**). Quantification shows that NLS-GFP loss rate became statistically different between the bleb and body in under one second (**Figure 2B-E**). The total NLS-GFP loss percentage did vary some between wild type, VPA, and *Lmnb1-/-*, but the bulk behavior was similar. Specifically, the NLS- GFP loss percentage of the nuclear body was < 10% over the 8 seconds tracked while the bleb showed a significantly higher 25-35% NLS-GFP loss. Micromanipulation point compression induced nuclear rupture clearly recapitulates a differential NLS-GFP loss rate of the nuclear bleb and body, again supporting that the nuclear bleb is a restricted compartment if the nucleus.

Non-bleb-based nuclear ruptures provide the ability to induce rupture via micromanipulation point compression and measure the loss dynamics in the same compartment. In non-bleb-based ruptures induced by moving point compression cause faster loss of NLS-GFP at a site of rupture compared to a location distal to the site of rupture for ∼2.5 seconds. Different from bleb-based ruptures, post 2.5 seconds the NLS-GFP loss percentage becomes similar for site of rupture and distal nuclear body site. Thus, the loss of NLS-GFP from the body can match that of the rupture site in a non-bleb scenario. Thus, it appears that non-bleb-based ruptures follow the same compartment dynamics on intervals > 2.5 seconds while bleb-based ruptures show signs of restricted compartmentalization due to differential dynamics lasting 10s of seconds between the nuclear bleb and body.

We also tracked the outcome of our micromanipulation point compression across conditions and nuclear shapes. Specifically, we show the percentage chance of bleb-based rupture (orange), bleb formation but no rupture (gray), no event (white), and non-bleb-based rupture (blue, **Figure 2F**). Interestingly, almost exclusively bleb-based nuclear rupture occurred only in nuclei already presenting a nuclear bleb in wild type, VPA-treated, and *Lmnb1-/-*. Abnormally shaped nuclei in *Lmna -/-* or normally shaped nuclei from all conditions almost exclusively resulted in non-bleb-based ruptures at varying frequencies upon micromanipulation moving point compression. The exclusion was normally shaped nuclei from VPA-treated cells which showed rare rupture of both types. Finally, we note that the majority normally shaped *Lmna-/-* nuclei under moving point compression resulted in non-bleb-based nuclear rupture confirming these nuclei are fragile. Overall, we find that micromanipulation moving point compression can induce rupture of already formed blebs but cannot form nuclear blebs leading to their rupture in normally shaped nuclei.

### FRAP reveals differential recovery dynamics of the nuclear bleb vs. body

Fluorescence recovery after photobleaching is established to determine turnover dynamics (Lippincott-Schwartz *et al*., 2018). We photobleached a portion of the nucleus and time lapse imaged at 5 second intervals to determine dynamic turnover of NLS-GFP in live cells. Photobleaching of a side of the nucleus led to near instantaneous recovery when measuring NLS- GFP signal intensity at the site of bleaching and distal to the site of bleaching (**Figure 3, A and C**). This agrees with moving point compression in non-bleb ruptures, which normalize loss dynamics in the same compartment in 2.5 seconds, or less than 5 seconds (**Figure 2F**). Thus, NLS-GFP turnover is faster than our ability to bleach and image inside of normally shaped nuclei.

**Figure 3.**
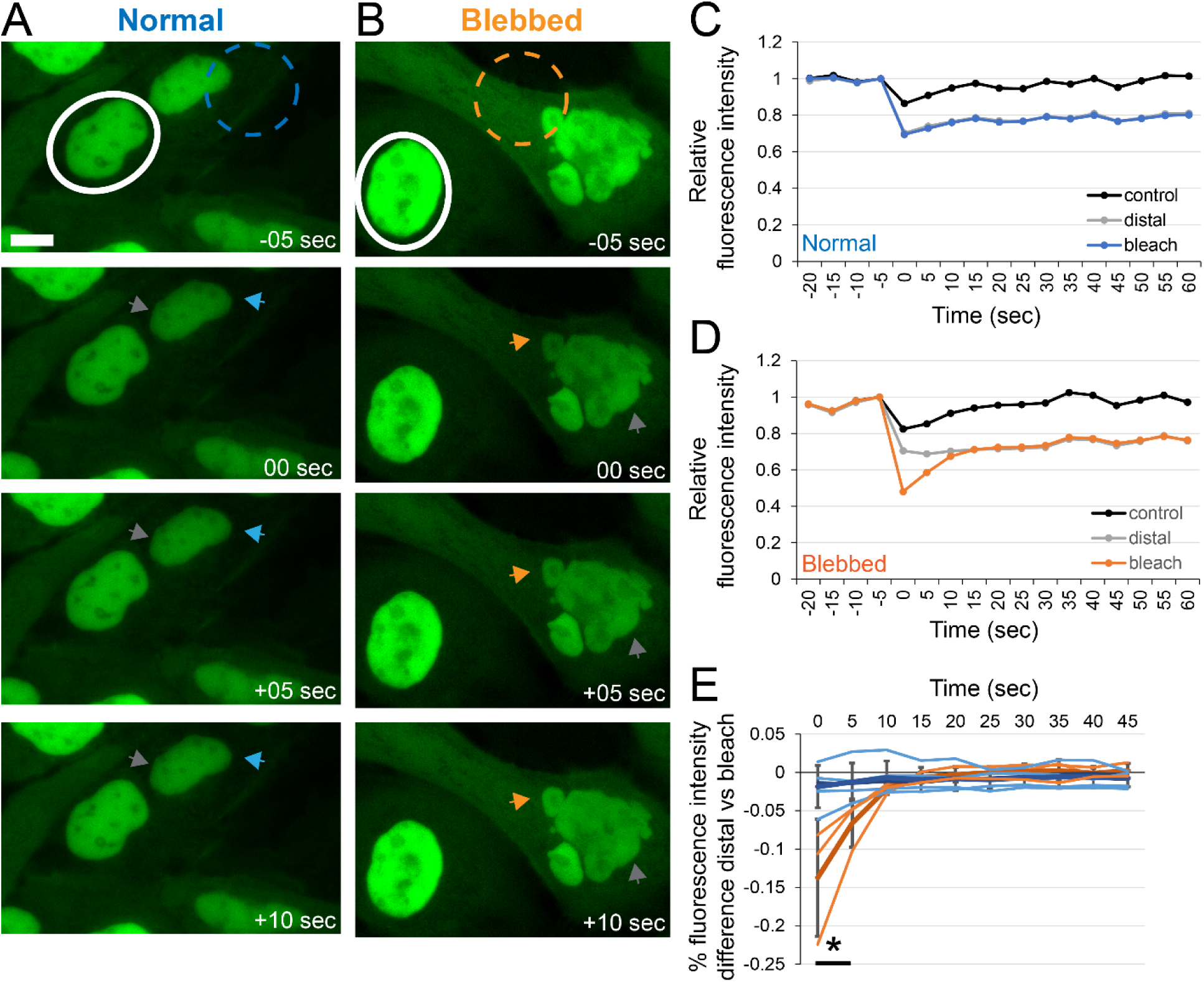
Fluorescence recovery after photobleaching supports nuclear blebs are restricted compartments in unruptured nuclei. (A, B) Representative images of fluorescence recovery after photobleaching (FRAP) of NLS-GFP in a (A) normal and (B) blebbed nucleus. Dashed line represents the bleach ROI for normal (blue) and blebbed (orange) while the white outline denotes the control/unbleached nucleus. Arrows track bleached area relative to gray arrows measuring the distal nuclear body location. Scaler bar = 10 µm. (C, D) Graphs of the representative FRAP event tracking control nucleus (black), bleach region (blue or orange), and distal part of the bleached nucleus (gray) for a (C) normally shaped nucleus and (D) the bleb of a blebbed nucleus. (E) Graph showing individual (lighter) and average (darker) of normal (blue, n = 4) and blebbed (orange, n = 3) nuclei. An asterisk denotes significantly different, p < 0.05. Error bars represent standard error.

We hypothesized that since the nuclear bleb was a restricted compartment of the nucleus FRAP of NLS-GFP in the nuclear bleb would take longer to recover than FRAP of a portion of a non- blebbed nucleus. Upon FARP in the nuclear bleb, NLS-GFP showed a delayed recovery on the order of 10-15 seconds (**Figure 3,B and D**). This delayed recovery of NLS-GFP into the nuclear bleb was statistically significant between the bleached bleb and distal nuclear body region as well as between bleb-based vs. normally shaped nucleus (**Figure 3E**). FRAP recovery of a U-shaped nucleus with a visible constriction showed similar loss and slow recovery but on a longer 30-60 sec time scale, likely due to its large size (**Supplement Figure 2A**). Micronuclei are separate compartments from the nuclear body (Krupina *et al*., 2021). FARP of a micronucleus proximal to a nuclear body showed slower and incomplete recovery of NLS-GFP signal (**Supplement Figure 2B**). Taken together FRAP data recapitulates differential dynamics between the nuclear bleb and body suggestive of the bleb being a restricted compartment from the nuclear body.

### 3D imaging reveals a significant constriction between the nuclear bleb and body

The differential dynamics of NLS-GFP movement between the nuclear bleb and body suggest that a constriction exists. Confocal imaging to provide nuclear size and shape of the nuclear body, bleb, and the transition between the two, we refer to as the neck, qualitatively shows a constricted neck between the nuclear bleb and body (**Figure 4A-C**). Measurements small width in the x and y-axis does not provide definitive data of a constriction. Instead, nuclear heights provide a clear measure for possible constriction in size between these three elements of the same nucleus including the bleb, neck, and body. Across multiple nuclei we find that the nuclear body is on average the largest height at 4.9 ± 0.2 µm while the nuclear bleb is smaller at 3.2 ± 0.2 µm. Measuring the transition between the nuclear bleb and body, we refer to as the neck, reveals a constriction of nuclear height to 2.3 ± 0.1 µm that is statistically smaller than both the bleb or body (**Figure 4, D and E**). Thus, there is a significant height constriction between the nuclear bleb and body that could cause disrupted diffusion dynamics of molecules in the nucleus.

**Figure 4.**
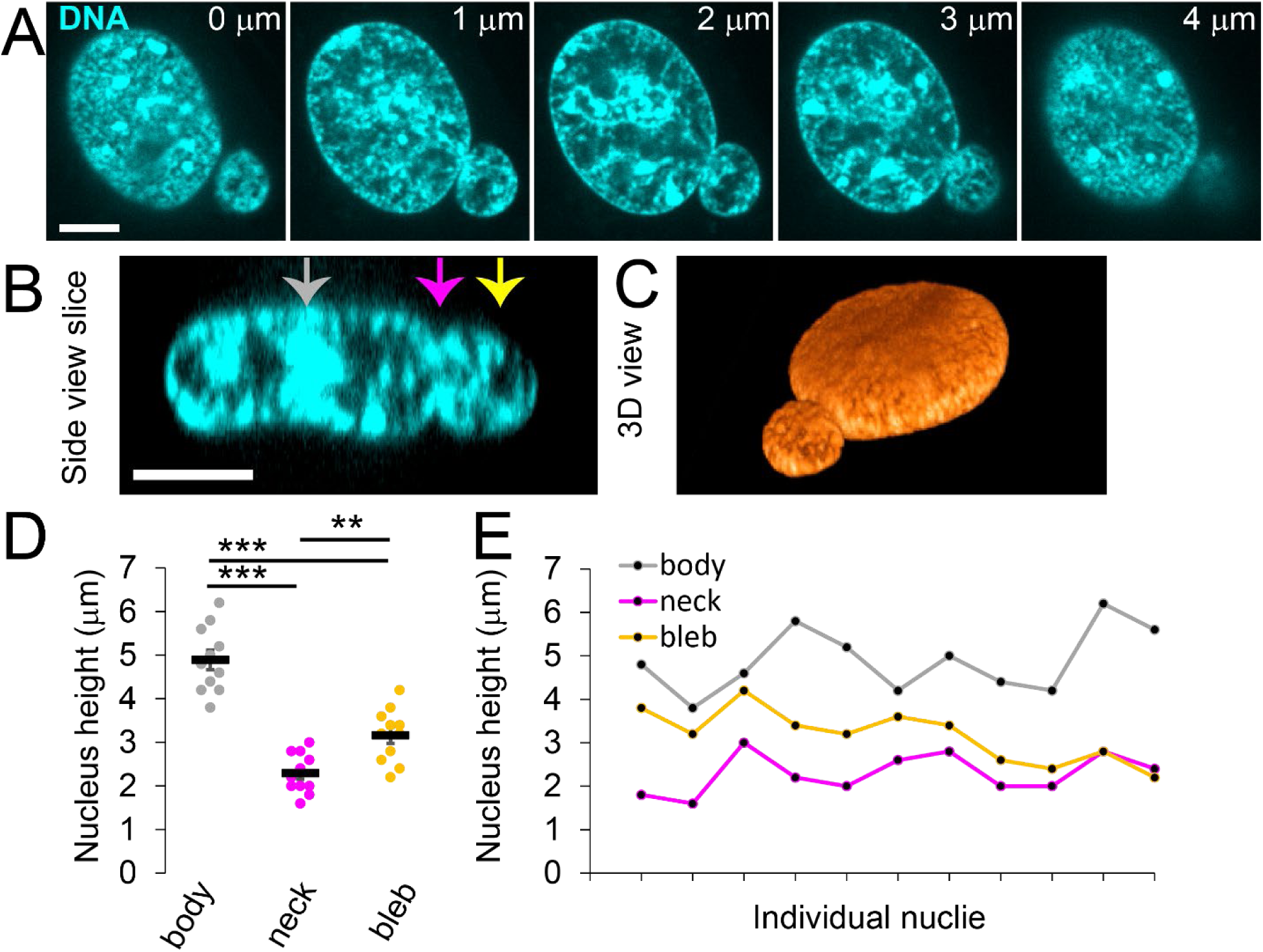
Height measurements confirm constriction between the nuclear body and bleb. Example confocal images of a Hoechst-stained blebbed nucleus (A) viewed top down (x,y) at 1 um focal plan increments, (B) side view slice denoting body (gray), neck (magenta), and bleb (yellow), and (C) 3D view. (D) Graph height of body (gray), neck (magenta), and bleb (yellow) measured by full-width half-maximum of fluorescence signal from side view. (E) Graph of each single nucleus body, neck, and bleb. Confocal images were taken at 200 nm z-steps. Statistical significance noted by asterisks ** p < 0.01 and ** p < 0.001. Scale bar = 10 µm.

## Discussion

Here we provided novel physics understanding the restricted compartmentalization of nuclear blebs as well as the nuclear rupture strength. Nuclear ruptures are commonly measured as loss of NLS-GFP signal from the nucleus and gain in the cytoplasm (Vargas *et al*., 2012; Robijns *et al*., 2016; Pho *et al*., 2023). However, measurements of loss of NLS-GFP from nuclear subsections or gain to cytoplasmic subsections were relatively unknown. Our data provides both the location and strength of rupture. Furthermore, we find that spill rates are significantly faster in the nuclear bleb compared to nuclear body revealing the nuclear bleb as a restricted compartment of the nucleus.

The data supports that nuclear ruptures are high pressure events. Specifically, compartmentalization loss of NLS-GFP from a bleb-based nuclear rupture is orders of magnitude faster rate, bleb or body, than loss of compartmentalization during nuclear envelope breakdown in mitosis (**Figure 1**). This suggests that nuclear rupture is a high-pressure single location breakage event where as nuclear envelope breakdown for mitosis occurs through disassembly of nuclear pores and it occurs in many locations (Güttinger *et al*., 2009). This idea of high pressure agrees with a study showing that 10-30 kPA of pressure is needed to cause 25-35% percentage loss of NLS-GFP loss (Zhang *et al*., 2019). This agrees with our measured 25-40% NLS-GFP loss nuclear ruptures. Many other labs confirm that nuclear ruptures result in this level of NLS-GFP loss (Denais *et al*., 2016; Young *et al*., 2020; Gunn *et al*., 2024).

The differential flow rate of measured NLS-GFP between the nuclear bleb and body provides non-mutually exclusive possibilities that the constrictions is large or that local pressure in the bleb is greater than in the body causing slower diffusion. This is likely due to la Place pressure, pressure in a highly curved surface that occurs in the nuclear bleb (Xia *et al*., 2018). Higher pressure in the nuclear bleb is supported by the faster loss rate compared to the nuclear body, even when the rupture occurs in the body (**Figures 1 and 2**). Our measurements of the constriction reveal a µm or larger constriction occurs qualitatively in the x,y dirrection while the constriction in the z is significant it is also relatively large at µm scale. Overall, our data supports that nuclear blebs restricted molecular movement is likely due to both pressure differential between nuclear bleb and body as well as statistically significant 3D constriction.

Nuclear blebs are restricted compartments that could disrupt normal nuclear functions. Our combined data across natural nuclear ruptures, induced nuclear ruptures via micromanipulation, and FRAP reveal differential dynamics between the nuclear bleb and body (**Figures 1-3**). Furthermore, we provide measurements showing a restriction at the neck between the nuclear bleb and body. The nuclear bleb as a restricted compartment might maintain or exclude molecules leading to dysfunction. The most commonly reported dysfunction upon nuclear rupture is increased DNA damage (Stephens, 2020; Kalukula *et al*., 2022). However, recent data has revealed that nuclear deformations (Pfeifer *et al*., 2018; Shah *et al*., 2021) and nuclear blebs (Chu *et al*., 2025) induce increased DNA damage independent of nuclear rupture. Our data show that nuclear blebs are restricted compartments provides a possible mechanism to increase DNA damage in nuclear blebs though inaccessibility to repair factors.

Transcription stalling is another mechanism that could occur in restricted compartment nuclear blebs. One key hypothesis is that actively transcribed genes end up in the bleb, but then become inactivated by being located in the nuclear bleb (De Vos *et al*., 2011; Helfand *et al*., 2012). It has been hypothesized this is could be due to the lack of RNA export due to loss of nuclear pores upon rupture (En and Ikegami, 2026). A non-mutually exclusive idea is that some degree of packing stabilizes transcription reactions (Miron *et al*., 2020; Paggi *et al*., 2026).

However, we have consistently shown that DNA density in the nuclear bleb is drastically decreased (Stephens *et al*., 2018; Bunner *et al*., 2024; Chu *et al*., 2025; Pujadas Liwag *et al*., 2025), and thus might be too low density for chemical reactions such as transcription and repair to work efficiently. We propose that actively stalled RNA Pol II likely in the initiating state in the nuclear bleb (Clark *et al*., 2026), causes open chromatin that is poised to cause translocations. The inability of open diffusion between nuclear bleb to the nuclear body and/or cytoplasm communication could cause transcriptional stalling, DNA damage, and resulting translocations that drive human disease state (Noe Gonzalez *et al*., 2021). This theory is partially supported by prostate cancer in which it is known that Androgen responsive genes end up often in the nuclear bleb (Helfand *et al*., 2012). Then translocations in active AR responsive genes are known to be associated with cancer severity due to translocations. We suggest this is due to being trapped in the nuclear bleb restricted compartment. Overall, we detail that nuclear blebs can cause dysfunction either through high pressure ruptures or through being a restricted compartment.

## Materials and Methods

### Cell culture and drug treatments

MEF WT, MEF *Lmnb1-/-*, and MEF *Lmna-/-*, were cultured as previously outlined (Pho *et al*., 2023) in DMEM (Corning) containing 10% fetal bovine serum (FBS, HyClone) and 1% penicillin/streptomycin (Corning). The cells were incubated at 37°C and 5% CO_2_, passaged every 2-3 days and kept for no more than 30 generations. MEFs were immortalized with SV40 large T antigen by retroviral transduction of the gene encoding the SV40 large T antigen as previously described (Shimi *et al*., 2011, 2015).

To treat with drugs, the cells were first plated in DMEM complete and incubated overnight. Cells were then treated overnight for 16-24 hours with 4 mM histone deacetylase inhibitor valproic acid (VPA, 1069-66-5, Sigma).

### Live cell NLS-GFP imaging and analysis

As previously outlined in (Pho *et al*., 2023), images were acquired with Nikon Elements software on a Nikon Instruments Ti2-E microscope with Crest V3 Spinning Disk Confocal, Orca Fusion Gen III camera, Lumencor Aura III light engine, TMC CLeanBench air table, with 40x air objective (N.A 0.75, W.D. 0.66, MRH00401). Live cell time lapse imaging was possible using Nikon Perfect Focus System and Okolab heat, humidity, and CO2 stage top incubator (H301). Cells were imaged in either 4 well cover glass dishes or 8 well cover glass chambers (Cellvis). For time lapse data, images were taken in 5-second intervals during 45 mintues of imaged 4 fields of view.

Images were saved within the NIS-Elements AR Analysis software. Images were observed to track nuclear blebs that resulted in nuclear rupture as well as non-blebbed nuclear ruptures in *Lmna-/-*. ROIs were drawn around the nuclear bleb and nuclear body to track NLS-GFP loss rates. ROIs were also drawn in the cytoplasm to measure cytoplasm/nucleus NSL-GFP intensity increase upon rupture. For tracking location of rupture 1 µm diameter circular ROIs were drawn in the cytoplasm around the nucleus. Data collected was then compiled and averaged in Excel (Microsoft) to determine percent blebbing, percent rupture, percent bleb-based rupture, and rupture frequency for each condition. Statistical significance was determined by conducting *t* tests between the two conditions.

### Live cell NLS-GFP imaging and analysis with micromanipulation

As previously described (Currey *et al*., 2022; Bunner *et al*., 2026), MEF NLS-GFP cells were grown in a micromanipulation well. Nikon Ts2R-FL microscope was used to image micromanipulation experiments. Using a pipette puller (P-97, Sutter Instrument, Table 1) a compression micropipette (TW100F-6) was tapered and then cut to 4 µm opening using MF-200 WPI microforge. This compression pipette was loaded into the IM-H1 Narishige holder and then the programmable micromanipulators (MP-285, Sutter Instrument). Using transmitted light phase imaging the compression micropipette was aligned with the cell nucleus before switching into green emission fluorescence imaging. Live cell fluorescence imaging was aquired at 30 frames per second as the compression micropipette was used to provide moving compression from the center of the nucleus to towards the periphery to induce nuclear blebbing and rupture. We calculated the overall strength of this moving compression as 10 nN using post-calibration to a known force micropipette.

Analysis was caried out in the same manner as naturally occurring nuclear ruptures outlined in the previous section.

### Fluorescence recover after photobleaching

MEF NLS-GFP cells were grown on 4 well cover glass dishes (Cellvis) for live cell imaging as outlined in the previous section “Live cell NLS-GFP imaging and analysis”. Photobleaching was accomplished by closing the CREST V3 IRIS to a 10 µm diameter circle and exciting with 364 nm light at full power for 1 second. Cells were imaged before and after photobleaching using Nikon Elements software at 5 second intervals. Analysis tracks the photobleached nucleus as well as a non-photobleached nucleus near by in the same field of view.

### Live cell imaging of nuclear height and analysis

Imaging and analysis were previously described in (Pho *et al*., 2023). Cells were grown in 4 well chamber glass dishes (Cellvis). Cells were stained with 1 µg/mL Hoechst 33342 (Life Technologies) for 15 minutes prior to imaging. Height images were taken on Nikon Eclipse Ti2 microscope with a 100x oil objective lens with Crest V3 Spinning Disk Confocal. Images were taken as 77 0.2 µm steps in 12 bit sensitive with exposure times between 50-100 ms. As described previously 2 intensity line scans were drawn approximately equal in distance from the nucleus center through the Z plane of each nuclei, values were exported to Excel, and the average of the full-width half-max of the two line scans was calculated.

## Acknowledgments

This work was primarily supported by NIH NIGMS grant Maximizing Investigators’ Research Award R35GM154928 to ADS. This work was further supported by NIDDK: 5K23DK144661 and NSF STC: 2535350 to LMA. NSF grant STC-2535350 and NIH grants U54CA268084, R01CA289294, R02CA225002, and philanthropic support from Susan Brice, the Christina Carinato Charitable Foundation, Rob and Kristin Goldman, and David Sachs to VB.

## Data availability

Raw data can be found here: https://doi.org/10.6084/m9.figshare.35016796.

## Author Contributions

Investigation, Methodology, Formal analysis, and Data curation were done by JC for nuclear rupture, MLC for micromanipulation moving point compression nuclear rupture, and MP for FRAP and nuclear height measures. KC, CH, and YB supported the paper through validation and formal analysis. VB and LMA aided supervision and writing through review & editing. ADS conceptualized the project, ran supervision, and was primarily responsible for original draft writing.

