## Supplemental Figures 1-2 for "Nuclear blebs are rupture-prone restricted compartments of the nucleus"

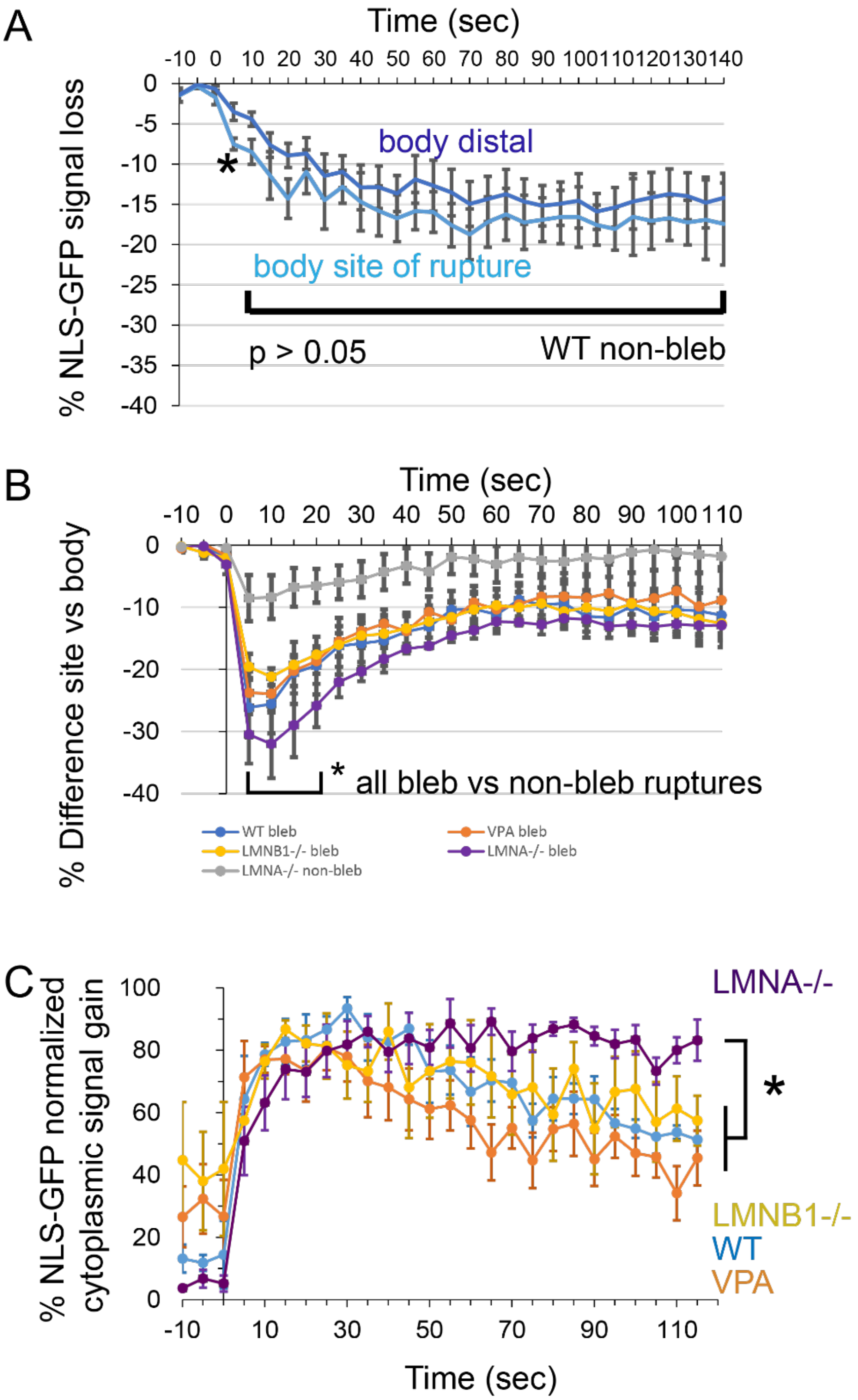


**Supplemental Figure 1. NLS percentage loss in the nucleus and gain in the cytoplasm.** (A) Average intensity of the main nucleus body (dark color) vs. body site of rupture for non-blebbed nuclear rupture for WT (n = 4). (B) Graph of the percentage NLS-GFP loss difference between the nuclear rupture site and the nuclear body for WT (B, blue, n = 5), VPA (C, orange, n = 15), *Lmnb1-/-* (D, yellow, n = 13), and *Lmna-/-* bleb (purple) and non-bleb (gray). (C) Graph of the percentage NLS-GFP gain in cytoplasm for WT blue, VPA orange, *Lmnb1-/-* yellow, and *Lmna-/-* purple.

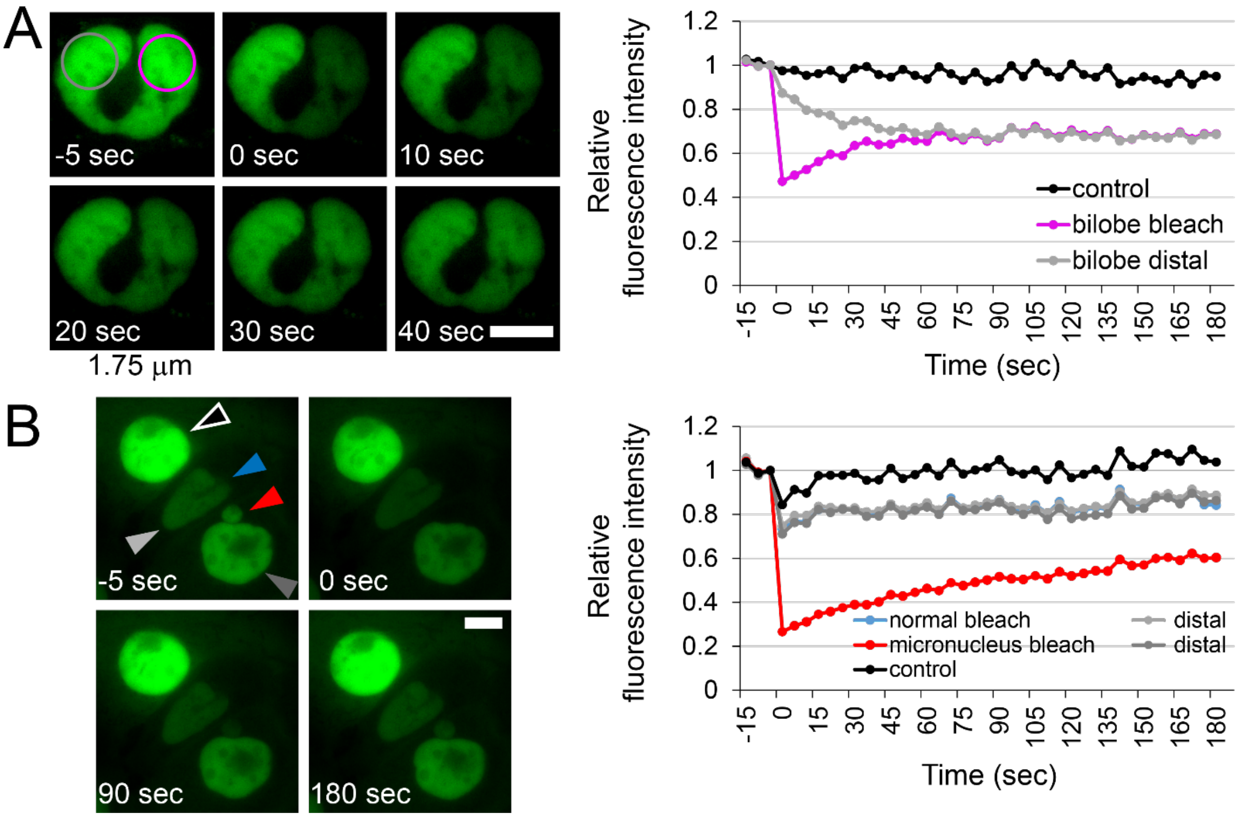


**Supplemental Figure 2. FRAP in bilobed nuclei and micronucleus.** (A) Example images and graph of a FRAP in a bilobed nucleus provides NLS-GFP dynamics through a 1.75 μm constriction as a proof of principal for restricted compartments. (B) Graph of FRAP relative fluorescence intensity changes for simultaneous bleaching of a separate component micronucleus (red) and a normally shaped nucleus (blue) each with their own distal non-bleached (gray). Scale bar is 10 μm.
